# STAMP: a multiplex sequencing method for simultaneous evaluation of mitochondrial DNA heteroplasmies and content

**DOI:** 10.1101/2020.07.05.188607

**Authors:** Xiaoxian Guo, Yiqin Wang, Ruoyu Zhang, Zhenglong Gu

**Author notes:** These authors contributed equally to this work (Xiaoxian Guo, Yiqin Wang). Regeneron Pharmaceuticals, Inc, Tarrytown, NY 10591, USA. Corresponding authors: Zhenglong Gu.

## Abstract

Human mitochondrial genome (mtDNA) variations, such as mtDNA heteroplasmies (the co-existence of mutated and wild-type mtDNA), have received increasing attention in recent years for their clinical relevance to numerous diseases. But large-scale population studies of mtDNA heteroplasmies have been lagging due to the lack of a labor- and cost-effective method. Here, we present a novel human mtDNA sequencing method called STAMP (<u>**s**</u>equencing by <u>**t**</u>argeted <u>**a**</u>mplification of <u>**m**</u>ultiplex <u>**p**</u>robes) for measuring mtDNA heteroplasmies and content in a streamlined workflow. We show that STAMP has high mapping rates to mtDNA, deep coverage of unique reads, and high tolerance to sequencing and PCR errors when applied to human samples. STAMP also has high sensitivity and low false positive rates in identifying artificial mtDNA variants at fractions as low as 0.5% in genomic DNA samples. We further extend STAMP, by including nuclear DNA-targeting probes, to enable assessment of relative mtDNA content in the same assay. The high cost-effectiveness of STAMP, along with the flexibility of using it for measuring various aspects of mtDNA variations, will accelerate the research of mtDNA heteroplasmies and content in large population cohorts, and in the context of human diseases and aging.

## INTRODUCTION

The human mitochondrial genome (mtDNA) is a 16.6kb circular genome encapsulated in the inner membrane of mitochondria. It encodes 22 tRNA and 2 rRNA genes used for mitochondrial protein synthesis as well as 13 evolutionarily conserved proteins in four of the five mitochondrial oxidation phosphorylation (OXPHOS) protein complexes (1). Although the primary and well-recognized function of mtDNA is encoding genes for energy production, a broad array of new biological processes and disorders associated with mtDNA have received increasing attention in the past decade (2, 3).

Recent deep sequencing studies have shown that mtDNA mutations are much more prevalent in human tissues than previously thought (4, 5). Given the multicopy nature of mtDNA in a single cell, mtDNA mutations can arise and co-exist with the wild-type allele in a state called heteroplasmy. mtDNA heteroplasmies can increase in fraction through clonal expansion in cells and tissues (6), without affecting mitochondrial function until their abundance reaches a certain threshold. At an intermediate fraction, a single disease-causing mtDNA mutation may lead to mitochondrial morphological changes and decreased transcription of mtDNA, recapitulating the mild mitochondrial dysfunction in diseases like diabetes and autism (7). At a relatively high fraction, it may induce global changes of gene expression involved in signal transduction, epigenomic regulation, and pathways implicated in neurodegenerative diseases (7). Accordingly, the varying fraction and abundance of mtDNA mutations, as well as their tissue sources, may give rise to distinct downstream phenotypes, which poses a challenge for mtDNA studies.

In addition to mtDNA sequence variations, the total number of mtDNA molecules in a cell, known as mtDNA copy number (i.e. mtDNA content), can also impact mitochondrial function (8). Altered mtDNA content in peripheral tissues is frequently reported in patients with neuropsychiatric disorders, and has been shown to be affected by stressful life events (9). Recently, several large-scale prospective studies found correlations between low mtDNA content in blood and age-related chronic diseases, such as cardiovascular diseases, illustrating that mtDNA content can serve as a biomarker for age-related decline of mitochondrial function, and a predictor for adverse health outcomes in humans (10, 11).

Lately, large-scale population studies on human mtDNA have been facilitated by widely available datasets from genome-wide sequencing projects. Off-target reads from whole-exome sequencing (WES) studies can be used to assess mtDNA sequence variations and medium-fraction heteroplasmies (12, 13). Likewise, whole-genome sequencing (WGS) with deep and uniform coverage on mtDNA can be used to measure mtDNA copy number and low-fraction heteroplasmies in tissues (14, 15). However, WES and WGS are not cost-effective for investigators to study mtDNA with a large sample size. Even if the genomic datasets have already been produced, analyses of mtDNA are often restricted by their original study design, which rarely allow investigators to study important characteristics of mtDNA, such as the temporal and tissue dynamics of mtDNA heteroplasmies and content.

mtDNA-targeted sequencing is an economical alternative to genome-wide methods. The main strategy is to isolate and enrich mtDNA from the total genomic background, and thus focus the sequencing capacity on mtDNA reads (16–18). These methods normally start with amplification of mtDNA from total genomic DNA (16, 17). Alternatively, mtDNA sequencing libraries can be enriched from total genomic sequencing libraries by using hybridization capture baits derived from mtDNA (18). Sequencing libraries containing short mtDNA fragments and adaptors are subsequently generated from the PCR products by using commercially available kits for sequencing library preparation, such as Nextra (16, 17) and QIAseq (19). However, as these library preparation protocols are optimized for processing large, linear genomic DNA, their use for the short 16.6kb circular mtDNA would dramatically increase the overall cost of mtDNA sequencing (20). To overcome this limitation, Nunez et al. developed a method to add sequencing adaptors directly to the short DNA fragments with T4 DNA ligase, after PCR amplification of mtDNA (21). Although this method can be applied to human mtDNA at a low cost, it has drawbacks that are similar to other methods, since mtDNA enrichment and library construction involve multiple steps and reaction plates, which incurs extra labor and increases the possibility of sample contamination during DNA purification and transferring.

Moreover, most of these mtDNA-enrichment strategies depend on high numbers of PCR cycles to increase mtDNA content, which inevitably introduce amplification errors and related mutation artifacts (22). These errors can lead to false discovery of mtDNA heteroplasmies, as most of them are of low fraction at a tissue level. A previous study showed that even at a sequencing coverage as high as 20000-fold (X) on mtDNA, the majority of the mtDNA sites were polymorphic with a variant allele fraction (VAF) of ≥0.1%, most of which could result from PCR or sequencing errors (16, 23).

Other methods that, conversely, reduce the level of the nuclear genome by using exonuclease V to digest the linear nuclear DNA (24), or by isolating mitochondria with differential centrifugation or magnetic beads for immunoprecipitation (25) require a large amount of starting materials, which is not suited for large-scale population studies with limited DNA sources or frozen tissue biospecimens. Importantly, by processing only mtDNA, the mtDNA-targeted sequencing methods lose valuable information on mtDNA levels in relation to nuclear DNA in the sample, making them unable to quantify mtDNA content in the same assay. Therefore, a cost-effective, accurate and flexible mtDNA sequencing method is urgently needed, especially for studying mtDNA variations in large populations.

In the current study, we developed a novel mtDNA-targeted sequencing approach called STAMP (<u>**s**</u>equencing by <u>**t**</u>argeted <u>**a**</u>mplification of <u>**m**</u>ultiplex <u>**p**</u>robes) which relies on multiplex single-stranded oligonucleotide probes to capture and subsequently sequence the entire 16.6kb human mtDNA. Furthermore, we extended STAMP to allow mtDNA content quantification in the same assay. Here, we discuss the design, workflow, cost-effectiveness and best practices of using STAMP for studying mtDNA heteroplasmies and content.

## MATERIALS AND METHODS

### Study samples

Two HapMap lymphoblast cell lines (sample 1: NA12751, and sample 2: NA18523) were purchased from Coriell institute. Upon receiving them, the lymphoblast cell lines were revived and cultured, at 37°C with 5% CO_2_, in RPMI 1640 medium containing 15% fetal bovine serum (VWR Life Science Seradigm, Inc.) and 1 × Antibiotic-Antimycotic (Thermo Fisher Scientific, Inc.). Total genomic DNA of these two samples was obtained using Wizard Genomic DNA Purification Kit (Promega, Inc.) as per the manufacturer’s instructions. The concentration of purified DNA was quantified by using a Qubit dsDNA HS assay kit (Thermo Fisher Scientific, Inc.). The five DNA sample mixtures were created by combining total genomic DNA of these two HapMap samples at relative ratios of 1:199, 1:99, 5:95, 20:80, and 50:50 (NA12751 versus NA18523). The lymphoblast cell line samples from 200 healthy individuals were collected in REGISTRY (26), and DNA were extracted as per REGISTRY protocol (26). Consent were obtained from all individuals enrolled in REGISTRY at site (26).

### mtDNA sequencing with STAMP

The experimental and computational protocols of STAMP used in the current study for sequencing mtDNA are listed as follows.

#### Step 1. mtDNA capture with extension-ligation (EL) probes

The oligos of EL probe pairs for each of the 46 mtDNA target regions and 5 nDNA target regions were column-synthesized, at 25 nanomole scale with standard desalting purification (Integrated DNA Technologies, Inc.). Equal amount of the 51 EL probe pairs were first pooled for mtDNA capture. In order to improve uniformity of sequencing coverage on mtDNA, several iterations from Step 1 to Step 5 were then performed to adjust EL probe pooling ratios. Hybridization reactions were performed on 50 ng genomic DNA with 4 ul EL probe mix and 1× Ampligase buffer in a 10 μl volume. Thermal conditions included 10 min at 95 °C for denaturation, followed by a decrease of 1°C per min to 55°C and 20h at 55°C for hybridization. 6 μl gap-filling mix including 0.1 mM dNTPs, 0.6M Betaine, 0.1M (NH_4_)_2_SO_4_, 0.5 units of *Tsp* DNA polymerase and 0.5 units of Ampligase in 1× Ampligase buffer was then added to the reaction mixture, which was incubated at 55°C for another 20 h for gap filling.

#### Step 2. PCR amplification of capture products

We used a dual indexing strategy to pool sample libraries for parallel sequencing (27). Each indexing primer comprised P5 or P7 Illumina adapter sequences, an 8-nt index sequence, a 13- or 14-nt pad sequence, and a universal sequence designed to anneal with the common termini of EL probes (**Figure 1B**) (27). PCR amplification was performed on 1.5 μl of capture product in a 50 μl PCR reaction with 1 × Phusion HF buffer, 0.5 μM of p5i5 and p7i7 indexing primers, 0.2 mM dNTP, and 1 unit of Phusion Hot-Start II DNA polymerase (Thermo Scientific, Inc.). PCR thermal conditions were 30 sec at 98°C for initial denaturation, followed by 25 cycles of 10 sec at 98°C, 15 sec at 65°C, and 15 sec at 72°C. The size and integrity of PCR products were visually verified by agarose gel electrophoresis.

**Figure 1.**
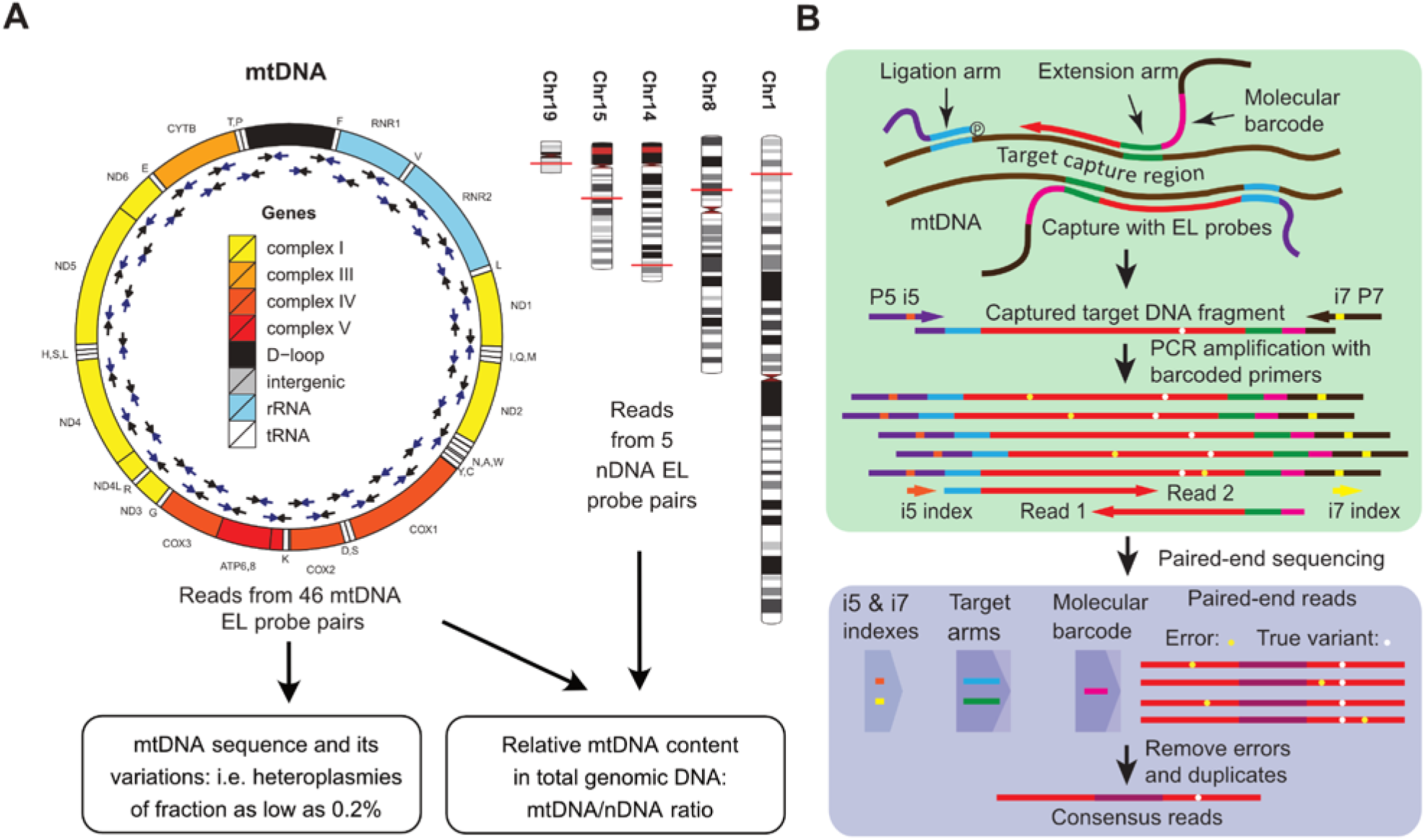
Design and workflow of STAMP. (**A)** Schematic diagrams of STAMP for mtDNA sequencing and relative mtDNA content assessment with EL probes. The locations of the 46 mtDNA EL probes are shown with pairs of arrows next to the mitochondrial genome. The locations of the 5 nDNA EL probes are shown with horizontal red lines across chromosomes 1, 8, 14 15 and 19. (**B**) Schematic diagrams for mtDNA capturing, gap-filling reaction, library construction, read processing, and consensus read calling in STAMP.

#### Step 3. Library purification

The obtained PCR products were purified and filtered by using AMPure XP magnetic beads with double size selection (Beckman Coulter, Inc.). In brief, 0.25 volume of beads was first used to bind DNA of >700bp in the PCR products, after which the supernatant was transferred to a fresh tube. An extra 0.4 volume of beads was added to bind DNA of >500bp in the supernatant. After the beads were washed and dried, DNA bound to these beads which contained PCR products of size in the range of 550bp to 650bp were eluted with 10 mM Tris-HCl, pH8.5, and were quantified with Qubit® 2.0 Fluorometer (Life Technologies, Inc.). Equal amounts of purified PCR products from different samples were pooled before parallel sequencing.

#### Step 4. Massively parallel sequencing

The sample libraries were sequenced with customized sequencing primers and 2×250 paired-end reads on Illumina sequencing flow cells. The Read 1 primer contained the13-nt pad sequence and the 20-nt universal sequence (TGCACGTCATCTACAGTAGGTCGGTGCGTAGGT), reverse complement of the sequence at the 3’ terminus of the ligation probe. The Read 2 primer contained the 14-nt pad sequence and the 20-nt universal sequence (CTCACTGGAGTTCAAGGGACGATGAGTGGCGATG) of the extension probe. The Index primer was the reverse complement of the Read 2 primer sequence (CATCGCCACTCATCGTCCCTTGAACTCCAGTGAG), which along with the complementary adapter sequences on the flow cell was used to read the dual sample indices (27). Cluster generation, image processing, and sequencing for samples of the current study were processed on MiSeq or HiSeq 2500 in the rapid run mode. Phi-X DNA library was spiked in at 5% to increase the complexity of the STAMP sequencing libraries.

#### Step 5. Sequencing data processing

A Python pipeline was developed to process and align paired-end reads generated from STAMP (**Supplementary Note B**). In brief, paired-end reads were first demultiplexed into files of individual samples based on the i5 and i7 index sequences. For each individual sample, paired-end reads were sorted into 51 clusters of capture products according to the arm region sequences identified at the locations of EL probes. The arm region sequences and the molecular barcode were trimmed from the paired-end reads, which were recorded in the read alignment files as annotations. To minimize complications from NUMTS in mtDNA read alignment, paired-end reads were first aligned to the reference human genome containing both nuclear DNA (genome assembly GRCh 38) and mtDNA (Revised Cambridge Reference Sequence, rCRS) sequences downloaded from ftp://ftp.1000genomes.ebi.ac.uk,using Burrows-Wheeler aligner (bwa mem, version 0.7.17) (28). Paired-end reads, annotated as having one of the 46 mtDNA EL probes, were marked as potential NUMTS in the alignment file if they could also be aligned to nuclear DNA with MAPQ≥10. Paired-end reads were aligned in a second round to a modified version of rCRS which had the final 120bp copied to the start to accommodate alignment of D-loop-region reads with the D10 probes. Paired-end reads that could not be aligned to the target region specified by their arm region sequences were removed. The remaining reads were locally realigned by using freebayes (version 1.1.0) (29) and their base qualities were subsequently recalibrated by using samtools (version 1.6) (30).

For paired-end reads with the same molecular barcode, the base information called at corresponding sites of the alignments was merged by using a Bayesian approach to generate a consensus read representing the captured mtDNA product. The same method was also used to merge base information within the overlapping region of the paired-end reads. The sequences of consensus reads were compared to a collection of known NUMTS sequences in the reference genome obtained from BLASTN search of the 46 mtDNA segments captured with EL probes, as well as their variant sequences harboring common polymorphisms (minor allele fraction > 1%) identified in the 1000 Genomes project (**Table S2**). A consensus read was marked as potential NUMTS if it showed a lower pairwise edit distance to NUMTS sequences than to the sample’s major mtDNA sequence, or if it was constructed from paired-end reads already annotated as NUMTS according to BWA alignment. Finally, consensus reads were converted to single-end reads, along with their base quality information, and stored in a bam file for each individual sample.

### mtDNA variant detection

mtDNA variants were determined by using consensus reads with MAPQ≥20 and BAQ≥30. Consensus reads marked as NUMTS or showing an excess of mismatches (>5 in the coding region and > 8 in the D-loop region; >11 for sample mixtures) compared to the individual’s major mtDNA sequence were also excluded from analysis. To reduce false positive calls of heteroplasmies, variants were subject to a list of quality filters, including (**1**) ≥100X depth of coverage with ≥70% of the bases having BAQ ≥30; (**2**) not in low-complexity regions (nt 302-316, nt 512-526, nt 16814-16193) or low-quality sites (nt 545, 16224, 16244, 16249, 16255, and 16263); (**3**) ≥5 minor alleles detected; (**4**) a log likelihood quality score of the variant ≥5; (**5**) comparable VAFs (Fisher’s exact test *P*≥10^−4^ and fold change ≤ 5) computed using consensus reads constructed with or without duplicate paired-end reads; (**6**) the detected number of minor alleles significantly larger than the expected number of errors, which was estimated at a rate of 0.02% in STAMP (Exact Poisson test, *P*<0.01/16569).

Functionalities for raw read processing, paired-end read alignment, consensus read calling, and variant detection have been implemented in the STAMP toolkit. The list of commands of the STAMP toolkit, arguments, and example code can be also found in **Supplementary note B**.

### mtDNA content evaluation with quantitative PCR

The relative mtDNA content of 126 lymphoblast samples, with enough genomic DNA after STAMP sequencing, was measured by using a quantitative PCR-based assay (31). The PCR reactions were performed as per manufacturer’s instructions (The Detroit R&D, Inc.). In brief, 15ng total genomic DNA was amplified with mtDNA or nDNA target primers and SYBR green PCR master mix in a 20 μL PCR reaction. Thermal conditions included 10 min at 95°C, followed by 40 cycles of 15 sec at 95°C and 60 sec at 60°C. For each sample, both mtDNA and nDNA targets were amplified twice in a total of 4 PCRs. Results from duplicates were averaged to compute mean Ct values for mtDNA and nDNA targets. The differences between them (ΔC_T_) were then normalized to that of a positive control sample measured on the same 96-well plate, by using ΔΔC_T_ method, to obtain qPCR-CN. qPCR-CN from 10 samples that failed in any of the 4 PCRs, and/or had a difference in CT values of over 3 cycles between experimental duplicates, were excluded from analysis.

### Statistical analysis

All statistical analyses were carried out with R (v3.5.0). The methods used and two-tailed *P* values are indicated in the main text.

## RESULTS AND DISCUSSION

### Design of STAMP

We designed single-stranded oligonucleotide probes to capture human mtDNA with an extension-ligation (EL) reaction (**Figure 1**). The extension probe has three parts, a 3’ extension arm with sequence complementary to the mtDNA target, a unique molecular tag used for tracking the capturing event, and a common PCR primer annealing region used for PCR amplification of the captured target. The ligation probe has a 5’-phosphorylated ligation arm with sequence complementary to the mtDNA target, along with another 20-nt common PCR primer annealing region at its 3’ end (**Figure 1B**).

To identify the required number of EL probe pairs and their mtDNA target locations, we first performed BLASTN search of human mtDNA sequences against the latest human reference genome (assembly GRCh38). We required that the resulting mtDNA segments be distinguishable from high-similarity segments derived from the human nuclear genome (**Tables S1 and S2**). Given the maximum sequencing read length of available Illumina sequencing platforms, we also required that the lengths of mtDNA targets should be around 400bp, so that they can be fully sequenced by using 2×250 or 2×300 paired-end reads, while the overlap between the paired-end reads is minimal.

As a result, we found that the entire 16.6kb human mitochondrial genome could be captured by using as low as 46 pairs of EL probes. We then placed the pairs of EL probes on the heavy and light strands of mtDNA alternatingly, to minimize the physical interference of adjacent probes in the multiplex reaction (**Figure 1A**). The locations and lengths of the extension and ligation arms in each of the EL probe pairs were further adjusted to ensure similar melting temperatures, around 55 °C, and similar GC-content, around 50%, and to avoid overlap with common mtDNA polymorphisms (population frequency > 1%) at the 3’ end of the extension arm and the phosphorylated end of the ligation arm (**Figure 1B**). In the end, we obtained 46 pairs of EL probes with a mtDNA target size ranging from 400 to 450bp to capture human mtDNA (**Table S1**).

### Effective capture of mtDNA with EL probes

In a pilot experiment, we synthesized the set of 46 pairs of EL probes (Integrated DNA Technologies, Inc.). We performed enzymatic gap-filling and ligation reactions on 50ng genomic DNA extracted from a lymphoblast cell line sample from the HapMap project, with 115 femtomoles of EL probe mixture. PCR amplification of the captured targets, using the 20-nt common PCR primers, requires the presence of PCR primer annealing regions at both ends, after successful polymerization of nucleic acids between the hybridized EL probes, as well as ligation of the polymerized nucleic acids with the 5’ end of the ligation arm. Therefore, captured products which lacked one of the common primer sequences, due to failed hybridization of either probe with its target sequence, or no ligation at the 5’ end of the ligation arm, could not be amplified.

Without adding genomic DNA templates or necessary enzymes, no clear bands were observed in the gel electrophoresis of PCR products from the gap-filling reactions (**Figure 2A**). In contrast, gel electrophoresis of PCR products from the effective gap-filling reaction on genomic DNA exhibited a smear, which had a size distribution centered at about 550bp-600bp, reflecting the expected sizes of PCR products comprising the target DNA, the molecular barcode, the probe arms, the common primers, and the sequencing adapters (**Figure 2A**).

**Figure 2.**
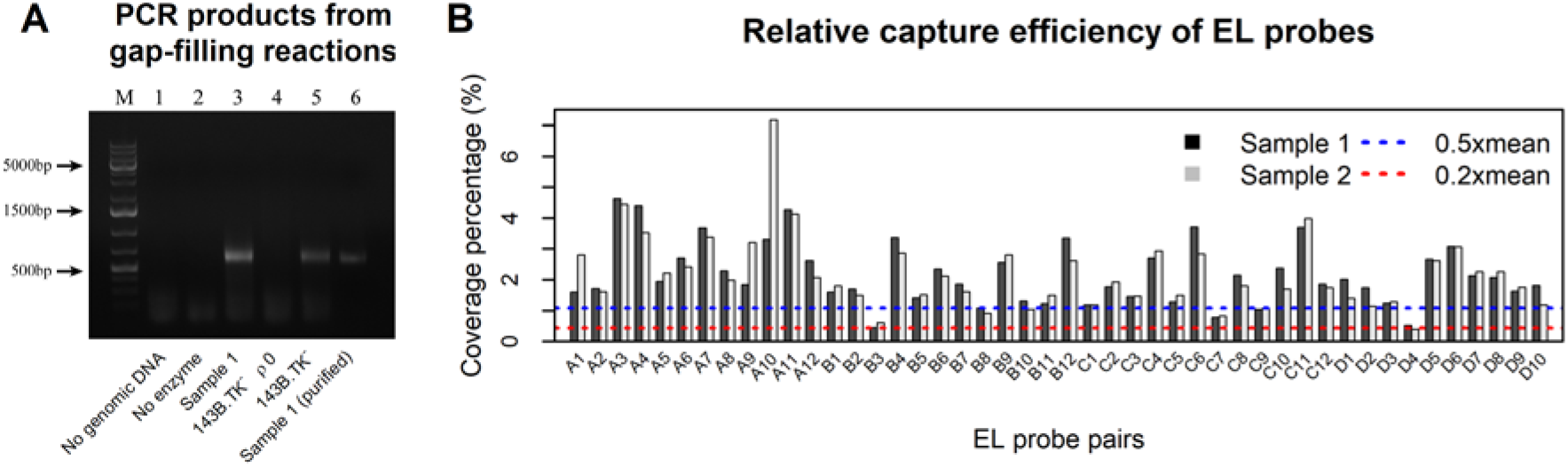
Effective capture of mtDNA with EL probes. **(A**) Agarose gel electrophoresis of the captured products: molecular-weight markers (lane M), no genomic DNA (lane 1), no gap-filling enzymes (lane 2), genomic DNA from a HapMap lymphoblast sample (lane 3), genomic DNA from 143B.TK^−^ ρ^0^ cell line (lane 4), genomic DNA from 143B.TK^−^ cell line (lane 5), and captured product after purification with double size selection (lane 6). (**B**) The relative depth of coverage of consensus reads on mtDNA for each of the 46 regions captured by EL probes (from A1 to D10 listed in Table S1). The blue dotted line and red dashed line indicate 50% and 20% of the mean sequence coverage, respectively.

Next, we purified PCR products with AMPure XP magnetic beads (Beckman Coulter, Inc.) and sequenced them with 2×250bp paired-end reads on MiSeq (Illumina, Inc.). After mapping the paired-end reads to the reference human genome, we found that all of the 46 mtDNA target regions were covered with reads at an average sequencing depth of 3512X, confirming that the set of 46 pairs of EL probes was able to capture the full length of human mtDNA (**Table S3** and **Figure 2B**). We further replicated these results in a second lymphoblast sample from the HapMap project (**Table S3** and **Figure 2B**). There was also a high correlation of coverage of regions captured by the same EL probe pair between these two samples (Pearson’s *r*= 0.8, *P*=3.3×10^−11^).

Moreover, capturing mtDNA sequences from genomic DNA could be potentially biased by the presence of nuclear DNA regions with high sequence similarity to mtDNA (i.e. nuclear mitochondrial segments, NUMTS) (32). To reveal its influence on mtDNA capture in STAMP, we performed gap-filling and ligation reactions with the set of 46 EL probes on genomic DNA from 143B.TK^−^ mtDNA-less (ρ0) cell line, and compared its PCR products with those obtained from the parental cell line 143B.TK^−^ which contains mtDNA. The gel electrophoresis of corresponding PCR products showed that the characteristic smear was only detectable in 143B.TK^−^ cells, but was absent in 143B.TK^−^ ρ0 cells, indicating that the resulting PCR products were largely derived from captured mtDNA target regions rather than off-target NUMTS (**Figure 2A**).

### Utilizing molecular barcodes to remove PCR duplicates and improve sequencing accuracy

Most mtDNA heteroplasmies are at low factions at the tissue level, which require a high depth of coverage of reads to reveal the presence of the variant allele and assess their fraction in relation to the wild-type allele. However, an ultra-deep read coverage (i.e. >2000X) on the small 16.6Kb human mtDNA may create errors in removing read duplicates if they are solely determined based on alignment coordinates and read sequences (33). PCR amplification of mtDNA before sequencing may also introduce biases in estimating VAF of a heteroplasmy (34). To resolve these issues, in STAMP, a 12- or 15-random-nucleotide molecular barcode was incorporated via the EL probe pairs to each of the capture products before PCR amplification, creating an identity for each capturing event (**Table S1**). Therefore, paired-end reads from the same mtDNA fragment captured in STAMP, including duplicates, can be determined according to the attached barcode information (**Figure 1B**).

Moreover, nucleotide mismatches at corresponding sites of paired-end reads with the same molecular barcode would suggest either PCR artifacts or sequencing errors (**Figure 1B**). We employed a Bayesian approach to merge the base information of paired-end reads with identical molecular barcode, generating a consensus read representing the captured DNA fragment (**Supplementary Note B**). We found that the number of nucleotide mismatches between the consensus read and the reference mtDNA sequence significantly decreased after merging base information of paired-end reads (Kolmogorov-Smirnov test, *P*<2.2×10^−16^, **Figure 3A**). Consensus reads with an excess of mismatches (number of mismatches, NM > 5) in comparison to the reference mtDNA were almost undetectable if they were constructed with duplicate paired-end reads (multiple reads with identical molecular barcode), which is 30-fold less frequent than those of consensus reads constructed from single paired-end read (Chi-squared test, *P*<2.2×10^−16^).

**Figure 3.**
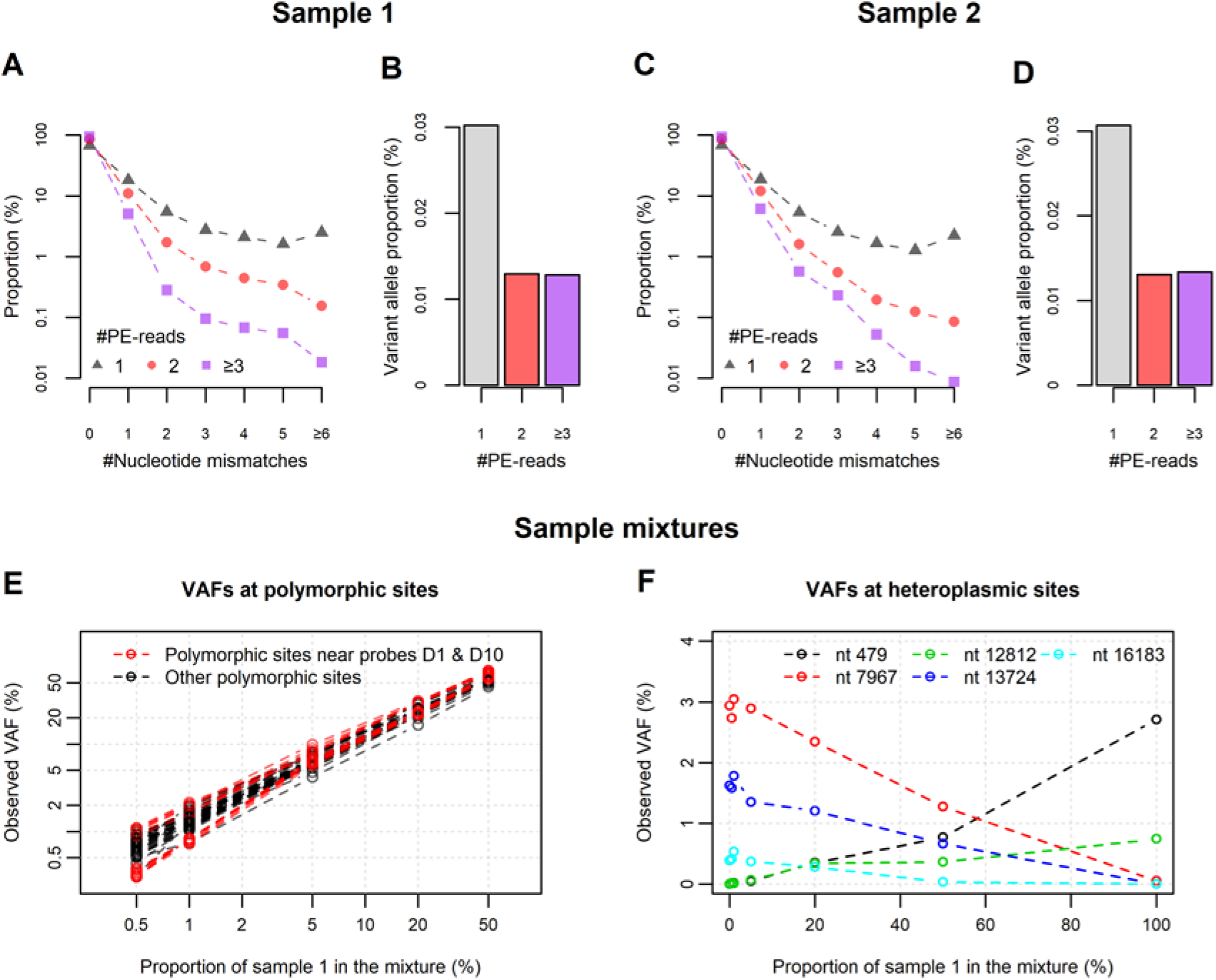
Accurate detection of mtDNA variants in sample mixtures. **(A, C)** The numbers of nucleotide mismatches between consensus reads and the reference mtDNA sequence in (**A**) sample 1 and (**C**) sample 2. **(B, D)** The proportions of variant alleles per base in the consensus reads after filtering out low-quality bases (BAQ<30) and consensus reads with excessive mismatches (NM>5) in (**B**) sample 1 and (**D**) sample 2. #PE-reads: the number of paired-end reads used to construct the consensus read. The VAFs of the mtDNA variants detected in the mixtures of sample 1 and sample 2 were depicted in **E** for variants at the 58 polymorphic sites and in **F** for variants at the 5 heteroplasmic sites. Each dotted line in **E** and **F** refers to the VAF changes of one variant in relation to the sample proportion indicated by the values on the x axis. Both x and y axes in **E** are shown on a log scale, and VAFs are shown in relation to mtDNA of sample 1,

After filtering out consensus reads with an excess of mismatches (NM>5) and bases with low quality scores (BAQ<30), we found that the proportion of variant alleles, which encompassed PCR and sequencing errors as well as low-level heteroplasmies, was about 0.013% and 0.03% per base among consensus reads with and without duplicate paired-end reads, respectively (**Figure 3B**). Both values are considerably lower than the reported proportions of ~0.1% per base in commonly-used mtDNA-targeted sequencing methods (35, 36). The proportion of variant alleles, at 0.013% per base, is also close to the error rate of *Tsp* DNA polymerase used in the gap-filling reaction of STAMP, which can be further improved using DNA polymerase with a higher fidelity (37). We obtained similar reductions in the distribution of nucleotide mismatches, and variant allele proportions, in the consensus reads of the second sample (**Figure 3 C, D**).

### Accurate detection of mtDNA heteroplasmies and their fractions in total genomic DNA

To evaluate the sensitivity and specificity of STAMP for detecting mtDNA heteroplasmies and their fractions in total genomic DNA, we applied STAMP to a series of sample mixtures created by combining total genomic DNA from the two lymphoblast samples used in the pilot experiment at varying ratios, ranging from 1:199 to 1:1 (**Table S3**). mtDNA sequences of these two lymphoblast samples differ at 59 single nucleotide sites (**Table S4**). One site (nt 16189) was in a low-complexity poly-C region of mtDNA and was excluded from the analysis of heteroplasmies. The average mean depth of coverage of consensus reads on mtDNA among the 5 sample mixtures was 3938X (median depth: 3284X), comparable to that of the two original samples at 3988X (median depth: 3392X; **Table S3**).

We found that all 58 polymorphic sites exhibited changes in their VAFs in accordance with the ratios of DNA used to generate these sample mixtures (*r*>0.997, *P*<0.00017; **Figure 3E** and **Table S4**). Of note, 2 polymorphic sites were located within the annealing regions of the EL probes: nt16519 at the 2^nd^ position from the 3’end of the extension arm of probe D10, and nt13650 at the 10^th^ position from the 3’end of the extension arm of probe D1. These two polymorphisms did not abolish the hybridization of EL probes to the target regions (for reasons discussed in the next section), but the variance in the VAFs of the 18 heteroplasmies close to the annealing regions of probes D1 and D10 was increased compared to that of the other heteroplasmies, especially in low-fraction sample mixtures (2-sided F-test *P*<1.4×10^−5^, **Figure 3E**). This suggests that a nucleotide mismatch between the arm regions of EL probes and their annealing regions in mtDNA can alter DNA capture efficiency, affecting the estimation of VAFs of nearby heteroplasmies in linkage with the nucleotide mismatch.

However, our pilot experiment using the sample mixtures represents an extreme scenario where all the 58 polymorphisms are in complete linkage with each other, and their alleles are separable into two haplogroups of mtDNA. In real human samples, the incidence of medium- and high-fraction heteroplasmies is usually low (38), and new heteroplasmies tend to arise in different mitochondria (39). Therefore, the variant and wild-type alleles of a heteroplasmy tend to share the same flanking mtDNA sequences. Both alleles would be captured by EL probe pairs at the same rate.

Next, we examined the influence of applying rigid quality control filters on detecting low-fraction mtDNA variants in the 3 sample mixtures, created with genomic DNA ratios of 1:199, 1:99 and 5:95. We found that 5 out of the 174 (3×58) variants were unable to survive the quality filtering procedures described in **Materials and Methods**. Of these, three showed a low percentage of high-quality reads and two were located at sites that did not have sufficient reads containing the variant alleles (**Table S5**). Moreover, all the 174 variants had VAF greater than 50% of the ratios of the genomic DNA in the sample mixtures (**Table S4)**. Therefore, STAMP’s sensitivity of identifying low-fraction heteroplasmies (VAF=0.5%-5%) is over 97%, with a cutoff for VAF at 0.25% and an average coverage of consensus reads at about 4000X.

By using these quality control filters, we found 17 other mtDNA heteroplasmies at VAF≥0.25% (**Table S5**). All of them were located at the 5 heteroplasmic sites already detected in one of the two original mtDNA samples, at a VAF from 0.4% to 2.9% (**Table S4**). These 5 heteroplasmic sites also displayed VAF changes proportional to the ratios of the DNA in the sample mixtures (*r*>0.9, *P*<0.0046; **Figure 3F**). Therefore, the false positive rate of STAMP in detecting heteroplasmies of VAF≥0.25% is under 10^−4^ (1/16569) per site of mtDNA.

### Application of STAMP in a population study of mtDNA heteroplasmies

To demonstrate the effectiveness and robustness of using STAMP for assessing mtDNA heteroplasmies in larger numbers of samples, we used STAMP to sequence mtDNA in 200 lymphoblast samples collected in REGISTRY (26). These 200 lymphoblast samples constituted the healthy control group of our research project on Huntington’s disease, and were sequenced on HiSeq2500 along with other samples relating to this project.

We were able to generate sequencing libraries for 192 (92%) out of 208 samples, including the experimental replicates from 8 lymphoblast samples. Among the 192 samples with STAMP libraries, 190 (99%) libraries from 182 lymphoblast samples were sequenced to >1000X depth of median coverage of consensus reads on mtDNA. The average median and mean depths of coverage of consensus reads on mtDNA were 4580X and 5450X, respectively (**Figure 4A**). The percentages of mtDNA sites that were covered with over 20% and 50% of the mean depth of coverage of consensus reads, as indicators for read coverage uniformity, were 98% and 78%, respectively (**Figure 4 B, C**). 99.4% mtDNA sites were covered with >500 consensus reads and 96.3% were covered with >1000 consensus reads. Similar to the observations from the two lymphoblast samples of the HapMap project, the overall proportions of variant alleles were 0.011% and 0.031% per base of consensus reads constructed with and without duplicate paired-end reads, respectively (**Figure 4D**). Taken together, these results confirm that STAMP can effectively sequence mtDNA in larger-scale studies.

**Figure 4.**
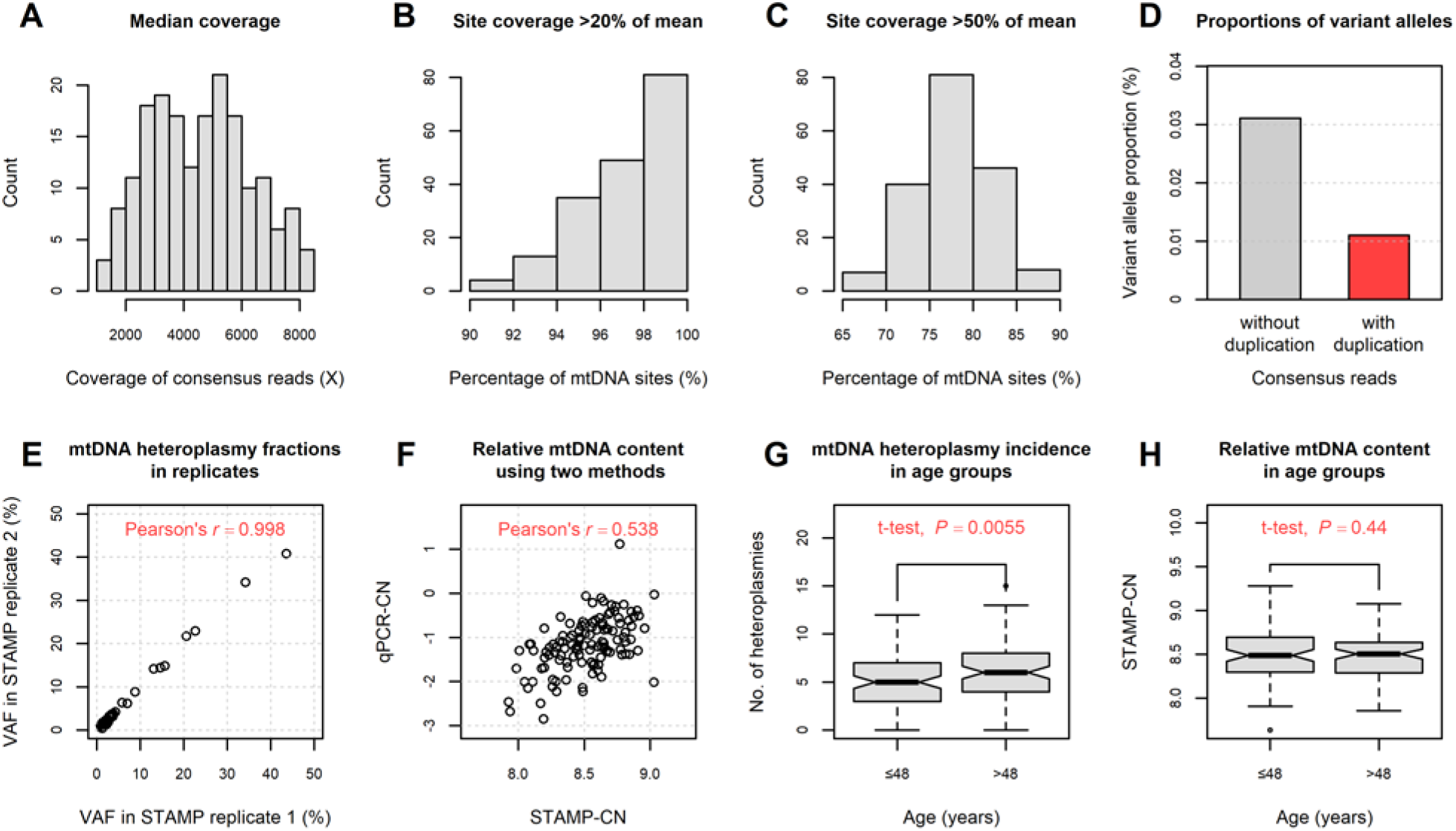
Reliable sequencing of mtDNA in a population study. Results of STAMP sequencing performed on 182 lymphoblast samples of REGISTRY are shown. (**A**) Median depth of coverage of consensus reads on mtDNA used for calling mtDNA variants. (**B** and **C**) Proportions of mtDNA sites with depths of consensus read coverage greater than (**B**) 0.2 and (**C**) 0.5 times the mean value, respectively. (**D**) Proportions of variant alleles per base in the consensus reads used for calling mtDNA variants. (**E**) Correlations of VAFs of 45 mtDNA heteroplasmies identified in the STAMP replicates performed on 8 samples. (**F**) Correlations of the relative mtDNA content measured by using STAMP and qPCR. (**G**) The box plots of mtDNA heteroplasmies detected in lymphoblasts of individuals aged above and under the sample median of 48 years old. (**H**) The box plots of the relative mtDNA content (STAMP-CN) detected in lymphoblasts of individuals aged above and under the sample median of 48 years old. VAFs in **E** were depicted using the major allele in replicate 1 as the reference allele.

Of note, two low-frequency mtDNA polymorphisms (nt2626 and nt15758) were identified at the 3’ end of EL probes A6 and D7 in two samples. These two mtDNA polymorphisms are single base transitions from A to G or T to C which give rise to purine-pyrimidine (A-C, C-A, G-T, and T-G) mismatches between the mtDNA templates and the arm regions of the EL probes. Since purine-pyrimidine mismatches in the 3’ end regions of primers have less detrimental impacts on PCR amplification than purine-purine or pyrimidine-pyrimidine mismatches (40), these two probes still produced 1248 and 268 consensus reads in the corresponding samples. These reads accounted for 0.3% and 0.4% of total consensus reads obtained, or equivalently 17% and 25% of the average proportions of consensus reads captured by EL probes A6 and D7 from mtDNA, respectively (**Figure S1**).

Since >90% of known polymorphisms, as well as heteroplasmies, in mtDNA cause single base transitions and are outside the 3’ end regions of EL probes (41), the current design of EL probes can be applied to capturing the entire length of most mtDNA haplogroups worldwide. However, any population other than Europeans, that were investigated in the current study, may lead to different coverage of consensus reads on mtDNA (**Figure S1**) if there are ethnicity-specific common polymorphisms or substitutions located in the arm regions of EL probes. Therefore, a pilot experiment of STAMP, aiming to identify appropriate molar ratios of EL probes that can ensure read uniformity on mtDNA for a population of interest, may be needed before large-scale applications. We further list all possible common polymorphisms (population frequency > 1%) located in the arm regions of EL probes in **Table S1** to help improve the design of EL probe sequences for non-European populations.

Among the 8 lymphoblast samples with replicated STAMP measures, we found that the major mtDNA sequences were identical between sequencing replicates. All the 45 heteroplasmies identified with VAF≥1% in one sample had VAF≥0.5% in the replicate, 44 (98%) of which also passed all quality control filters for heteroplasmy calling (**Table S6**). Overall, the correlation between VAFs of the heteroplasmies detected in sequencing replicates was estimated to be *r*=0.998 (*P*=3×10^−53^, **Figure 4E**). The average coefficient of variation in repeated measures of VAFs, at median site coverage of 4330X, was 5% for medium-to-high-fraction heteroplasmies (VAF≥5%; median VAF=15%) and was 17% for low-fraction heteroplasmies (0.5%≤VAF<5%; median VAF=1.4%). These values are close to the corresponding estimates of 4% and 13% computed using the sampling distribution of sample proportions at the VAF medians. These results indicate that STAMP can reliably detect heteroplasmies and quantify their fractions in genomic DNA.

### Extension of STAMP to measure mtDNA content in the same assay

We further explored the possibility of modifying STAMP to enable mtDNA content quantification in the same assay. To this end, we added five pairs of EL probes to capture single-copy regions in nuclear DNA (nDNA), along with the 46 pairs of mtDNA EL probes in STAMP. These five target nDNA regions are located on different autosomal chromosomes (**Figure 1A**). Assuming that the amount of DNA captured by each pair of EL probes is proportional to the number of copies of the target region in the human genome, reads from the nDNA regions then can be used as a normalization factor to adjust differences in total genomic DNA input and sequencing coverage across samples.

We first evaluated the performance of the nDNA EL probe pairs in capturing their target regions relative to the mtDNA EL probe pairs. We have noted in the previous analyses that the presence of polymorphisms in the arm regions of the EL probes could influence the capture efficiency of the target region. We thus focused on a subset of the 46 EL probe pairs to compute an average number of consensus reads for mtDNA. This subset comprised 18 EL probe pairs (A5-A8, B2, B6, B7, B9, C1-C5, C7, C9, C12, D1, D5) that lack common polymorphisms in their arm regions in European populations, and showed relatively low variations in consensus read coverage across samples of the current study (**Table S1** and **Figure S1**).

We found that all the five nDNA probe pairs exhibited positive correlations in their consensus read numbers with that of mtDNA EL probe pairs (R^2^=0.4-0.79, **Figure S2A-E**). To improve reliability in estimating nDNA content in the sample, we used the 3 EL probe pairs targeting chromosomes 8, 14, and 19 with an R^2^≥0.74 to compute an average number of consensus reads for nDNA. In addition, we found that the performance of EL probe pairs was not equal when capturing nDNA compared to mtDNA target regions (**Figure S2A-E**), possibly due to a compact design of EL probes on mtDNA. To adjust for this difference, we computed the relative mtDNA content for STAMP (hereafter referred to as STAMP-CN) as the average consensus read number from mtDNA relative to that from nDNA, by using the equation: log_2_(No. of mtDNA consensus reads)–C×log_2_(No. of nDNA consensus reads). C in the equation stands for the normalization factor for nDNA consensus reads, estimated using the coefficient β from the regression of log_2_(No. of mtDNA consensus reads) against log_2_(No. of nDNA consensus reads), which was equal to 0.53 (**Figure S2F**).

By comparing STAMP-CN with the relative mtDNA content (hereafter referred to as qPCR-CN) determined using a commercially available quantitative PCR-based assay performed on the same sample (42), we found a significant positive correlation (*r*=0.54, *P*=10^−5^, **Figure 4F**) between the values of STAMP-CN and qPCR-CN. This correlation is in good agreement with the result reported in a previous comparative study of mtDNA content measured by sequencing-based methods and qPCR-based methods (42). Because STAMP estimates the amount of mtDNA by directly counting reads from multiple regions in nDNA and mtDNA, STAMP-CN may be more accurate in reflecting real mtDNA content than that quantified based on fluorescence intensity from qPCR assays targeting single nDNA and/or mtDNA regions.

### Age-dependent increase of heteroplasmy incidence in lymphoblast samples

With both mtDNA heteroplasmies and mtDNA content measured in the same sample, we then evaluated how age impacts these mtDNA characteristics in lymphoblast samples as compared to what we previously observed in mtDNA of blood samples, using the WGS dataset of the UK10K project (14).

We identified 1007 heteroplasmies of VAF≥1% across the entire length of mtDNA in 182 lymphoblast samples of REGISTRY (**Figure S3A**). The average number of heteroplasmies per sample was 5.5 (range:0-15). 180 (99%) lymphoblasts possessed at least one heteroplasmy in mtDNA (**Figure S3B**). Similar to our previous study on lymphoblast mtDNA (5), the number of mtDNA heteroplasmies identified in lymphoblasts was consistently greater than those of whole blood, at about 1 heteroplasmy of VAF≥1-2% (14, 16), implying that mtDNA of lymphoblasts may enrich for pre-existing variants in somatic cells that are undetectable at a tissue level.

At the variant level, 760 (88%) of the 862 identified heteroplasmic sites were unique to one of the 182 lymphoblast samples **Figure S3C)**. Over half (54%) of the heteroplasmic sites did not overlap with known mtDNA polymorphisms (a population frequency < 0.01%), and another 20% were found to overlap only with rare polymorphisms in less than 0.1% of the general population (**Figure S3D**). The base changes of heteroplasmies showed a high transition to transversion ratio at 15. This suggests that the dominant mutational force underlying heteroplasmies is nucleotide misincorporation by polymerase gamma or deamination of bases in mtDNA, consistent with mtDNA mutation patterns identified in blood (43).

We first performed Student’s t-test to compare mtDNA heteroplasmies and content between individuals aged above and under the sample median of 48 years old. We found increased mtDNA heteroplasmy incidence and decreased mtDNA content in lymphoblast samples in the older group (mean age: 55 years old) compared to those in the younger group (mean age: 41 years old). But only mtDNA heteroplasmy incidence showed a significant difference between these two age groups (**Figure 4G**; Cohen’s d=0.42, *P*=0.0055). The lack of a significant association of mtDNA content with age (**Figure 4H**; Cohen’s d=-0.12, *P*=0.44 for STAMP-CN; Cohen’s d=-0.07, *P*=0.71 for qPCR-CN) might be due to insufficient statistical power, since only a mild annual decrease of 0.4 mtDNA copies (0.24% of the population average in 1511 individuals, *P*=0.0097) in blood was noted in our previous study (14). Furthermore, cell culture condition might affect mtDNA content (44), which overwrites the real association between age and mtDNA copy number that was observed in blood.

By using a *Poisson* regression model in the analyses of heteroplasmy incidence, we found an annual rate of increase of 1.2% (95% CI: 0.5%-2.0%; *P*=0.00063) for heteroplasmies of VAF≥1% in lymphoblast samples (**Table 1**, **model 1**). In line with our previous findings in blood (14). the increase of heteroplasmy incidence in lymphoblast samples with age was not affected by changes in mtDNA content (**Table 1**, **model 2**). Moreover, we found a similar age-dependent increase of heteroplasmy incidence after focusing on unique variants detected in our dataset, meaning that random genetic events in mtDNA, such as replication errors or drift, are largely responsible for the accumulation of mtDNA heteroplasmies during aging (**Table 1**, **model 3**). Significant age effects were also obtained using heteroplasmies of higher VAFs (VAF≥2% or VAF≥5%, **Table 1**).

**Table 1.** Age-dependent increase of mtDNA heteroplasmy incidence in lymphoblast samples.

| mtDNA heteroplasmy | model 1 |  | model 2 |  | model 3 |  |
| --- | --- | --- | --- | --- | --- | --- |
|  | Beta [95% CI] | *P* | Beta [95% CI] | *P* | Beta [95% CI] | *P* |
| **VAF ≥ 1%** | 0.012<br>[0.005-0.020] | 0.00063 | 0.012<br>[0.005-0.019] | 0.00073 | 0.014<br>[0.005-0.022] | 0.0012 |
| **VAF ≥ 2%** | 0.017<br>[0.007-0.027] | 0.00087 | 0.017<br>[0.007-0.026] | 0.00091 | 0.020<br>[0.009-0.031] | 0.00043 |
| **VAF ≥ 5%** | 0.027<br>[0.013-0.042] | 0.00025 | 0.027<br>[0.013-0.042] | 0.00026 | 0.037<br>[0.020-0.054] | $2.7 \times 10^{-5}$ |

The associations between heteroplasmy incidence and age were computed by using *Poisson* log-linear model with adjustment for sex and sequencing coverage in model 1, and for sex, sequencing coverage, and the relative mtDNA content (STAMP-CN) in model 2 and model 3. In model 3, only heteroplasmies that occurred once in the 182 lymphoblast samples were used to compute the incidence.

The observed rate of increase of heteroplasmy incidence in lymphoblast samples is in line with the estimate in our previous study of blood, showing a 59% increase in the heteroplasmy incidence between individuals in the youngest and oldest age groups, spanning an average of 44 years (14). These results indicate that the age-dependent accumulation of mtDNA heteroplasmies may be conserved in lymphoblast samples, after immortalization of B lymphocytes by Epstein-Barr virus and short cultivation of the cell lines. Therefore, lymphoblast samples may serve as a useful genetic resource for studying age-related mtDNA mutation spectra in the hematopoietic system, and their contributions to mitochondrial dysfunction in diseases associated with aging.

## CONCLUSION

In the current study, we present a novel human mtDNA targeted sequencing method, STAMP, which enables simultaneous assessment of mtDNA sequence variations and mtDNA content. Our method streamlines the experimental workflow with multiplex capture of human mtDNA and nDNA, and generates high-quality sequencer-ready libraries. This novel methodology eliminates the error-prone steps of transferring reagents and DNA samples, reduces the risk of DNA contamination, and enables mtDNA sequencing in thousands of samples.

Importantly, with high cost-effectiveness (**Figure S4** and **Table S7**) and a flexible design, STAMP can be used to study mtDNA variations at different scales and to determine mtDNA heteroplasmies at different fraction levels (**Table S8**; discussed in **Supplementary Note A**). Given the 0.01%-0.03% error rates of STAMP (**Figure 2 B** **and** **D**), STAMP can be used to detect heteroplasmies of fractions as low as <0.5%, with deeper sequencing coverage (**Figure S5**). Thus, STAMP can be used in studies of somatic mtDNA mutations in tissue specimens, which is currently unachievable by using other mtDNA-targeted sequencing methods, or whose experimental cost is prohibitive, when a large number of samples need to be sequenced.

Furthermore, we provide in the current study the related experimental details and computational solutions to assist the application of STAMP in future human mtDNA studies. Accordingly, the insights gained from these studies will transform our understanding of the role of mtDNA in aging and age-related diseases of humans.

## DATA AVAILABILITY

The STAMP toolkit used for processing sequencing reads from STAMP will be available for download at https://github.com/mtstamp/stamp.

## SUPPLEMENTARY DATA

Data in the supplementary PDF file consist of 2 supplementary notes (**Supplementary Note A** and **B**) and 5 supplementary figures (**Figures S1-S5**).

Data in the supplementary Excel file consist of 8 supplementary tables (**Tables S1-S8**).

## ACKNOWLEDGEMENTS

We thank NIH (R01AI085286), the CHDI Foundation and the European Huntington’s Disease Network for their support in this research project. We thank Andrew Cheung for assisting in mtDNA sequencing experiments. We also thank Drs. Chunyu Liu, Qi Sun, Paul Soloway, Haiyuan Yu, and Yiping Wang for critical reading and comments on the manuscript.

## AUTHOR CONTRITUTION

XG and ZG designed STAMP with inputs from YW and RZ. XG performed mtDNA sequencing experiments and YW provided computational tools. YW, XG and ZG analyzed the data and wrote the paper with inputs from RZ. All authors read and approved the final manuscript.

