## Supplementary Text for "STAMP: a multiplex sequencing method for simultaneous evaluation of mitochondrial DNA heteroplasmies and content"

#Current address: Regeneron Pharmaceuticals, Inc, Tarrytown, NY 10591, USA

### Supplementary Material

**Supplementary Note**

**A. Cost-effectiveness and flexibility of STAMP applications**

The integration of mtDNA capture and enrichment with multiplex probes in STAMP can effectively reduce the cost of sequencing library construction to be under $5 (**Table S7**). When we call mtDNA variants, the minimum VAF of the heteroplasmies and the statistical power to distinguish them from sequencing and PCR errors are both affected by read depths and error rates of sequencing. Both parameters can be adjusted in STAMP by changing the numbers of consensus reads and paired-end reads, allowing the sequencing costs and scales to be flexible according to the aim of the study (**Table S8**).

The number of consensus reads obtained for mtDNA in STAMP reflects the number of mtDNA fragments (NF) captured with EL probes. By fitting a Poisson distribution to the numbers of paired-end reads used in constructing consensus reads in the 182 lymphoblast samples of the current study (**Figure S4 C and D**), we found that NF was close to 6000 per EL probe target region in 1.5 ul of capture product from STAMP sequencing performed on 50ng of lymphoblast DNA. Yet, NF may vary, depending on the extraction methods, tissue sources and quality of the genomic DNA. Therefore, we recommend conducting a pilot experiment with STAMP on 10-20 samples in one lane of MiSeq to estimate NF empirically. The obtained NF can then be used to calculate the amount of capture product that needs to be amplified and sequenced, to ensure enough consensus reads for detecting mtDNA heteroplasmies.

For example, in the current study, 1.5 ul of capture product contained roughly an average of 6000 unique mtDNA fragment for each of the 46 EL probes. Among the 190 lymphoblast samples, the rate of paired-end reads retained for constructing consensus reads for mtDNA and nDNA was 0.9 and 0.003, respectively, after alignment and quality filtering (**Figure S4 A and B**). Given a yield of 125 million 2x250bp paired-end reads from one lane of a flow cell processed on HiSeq 2500, a batch load of 250 sample libraries on each lane can produce an average of about 10000 ($0.9\times125\times10^6/46/250$) and 300 ($0.003\times110\times10^6/5/250$) paired-end reads per EL probe region in mtDNA and nDNA, respectively, for each sample (**Table S8**). Accordingly, each consensus read will be constructed from an average of about 2 paired-end reads. About 60% of consensus reads will have duplication, which improves the error rate of STAMP from 0.03% to 0.02% per base (**Figure S5C**). At this error rate, STAMP guarantees >99% power to distinguish heteroplasmies of VAFs at 1% and 0.5% from errors, at an average of 98% and 78% of mtDNA sites, respectively (**Figure S5D**).

Similarly, very-low-fraction heteroplasmies can be detected by further increasing the numbers of consensus reads and paired-end reads. For example, 20,000 consensus reads per EL probe region and 80,000 paired-end reads can be achieved by amplifying 5ul of capture products, and sequencing the resulting libraries in a batch load of 31 samples on one lane of HiSeq 2500 (**Table S8**). As a result, >92.5% consensus reads will incorporate information from at least 2 paired-end reads, and, on average, 4 paired-end reads which lowers the error rate to 0.012% per base and provides >99% and >94% power for detecting heteroplasmies at VAF of 0.2% for 78% and 98% of mtDNA sites, respectively (**Figure S5 E and F**).

However, if the aim of the study is to assess medium- or high-fraction heteroplasmies, polymorphisms, or haplogroups, which do not require ultra-deep read depths to detect, increasing the number of either consensus reads or paired-end reads may waste sequencing capacity. Under these circumstances, a batch load of up to 1000 sample libraries per lane on HiSeq 2500 can be applied to achieving an average coverage of consensus reads and paired-end reads at 2000X and 2500X, which can be used to detect heteroplasmies of VAF ≥2% (**Table S8; Figure S5 A and B**). Moreover, according to the observed ratio of nDNA and mtDNA reads in the current study (**Figure S4 A and B**), an average of over 50 consensus reads can still be attained from the nDNA regions for computing STAMP-CN.

#### B. Implementation of the STAMP toolkit

We developed a python pipeline (the STAMP toolkit) to process sequencing data. Each functionality described in the main text has been implemented in the STAMP toolkit (hereafter referred to as stamp) and is summarized in the flow chart below. stamp has four modules, “align”, “pileup”, “scan”, and “annot”, which are described in the following sections.

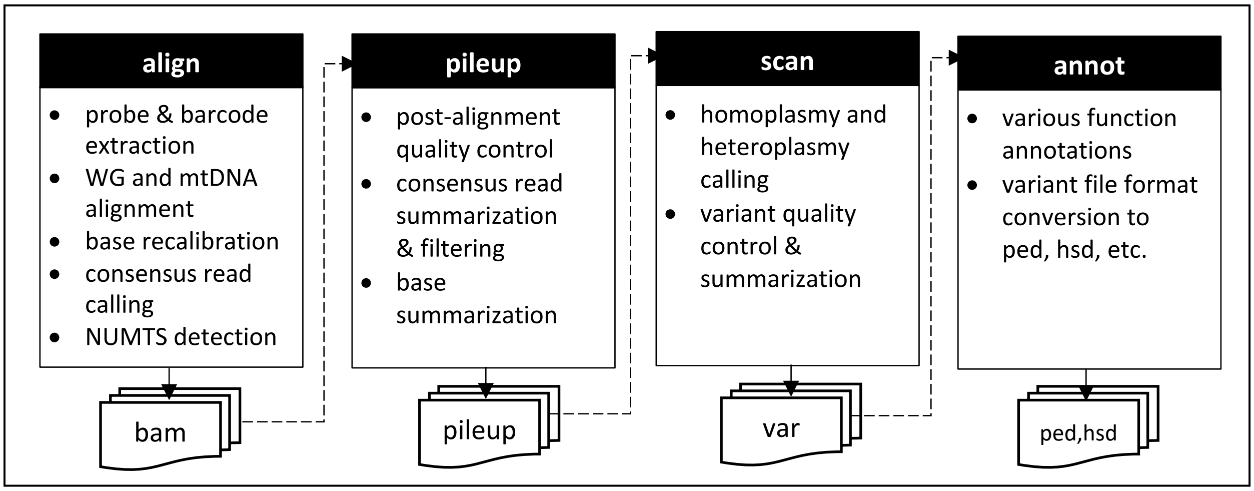


**Here is the command to list all the stamp modules:**

```
python stamp.py -h

    Usage: stamp.py <command> [options]
    Command: align     generate the consensus read alignments
             pileup    summarize the consensus read bases
             scan      variant identification
             annot     variant annotation
```

#### B1. Read alignment and consensus read calling

In the “align” module, stamp reads the raw fastq files, and extracts the probe arm and molecular barcode sequences from the paired-end reads according to the design of EL probes in STAMP (**Figure 1B**). The sequences of the probe arms must be from one of the 46 mtDNA and 5 nDNA probe pairs. Because of sequencing errors, a maximum of 3 nucleotide mismatches is allowed between the arm sequence and the matched probe sequence. Due to decreased quality scores at the start of the paired-reads, we only use 13 bases starting from the $3^{rd}$ position in the 15-nt molecular barcode for recording the capturing event, with a total of 67 million unique base combinations. This number is significantly greater than the depth of coverage of consensus reads observed in the current study and those listed in **Table S8**. The molecular barcode must also contain at least 9 bases with BAQ≥15. To further minimize the influence of sequencing errors on barcode sequences, we mask low-quality bases (BAQ<15) with "N" and do not count these bases in comparing barcode sequences between two paired-end reads.

The paired-end reads that pass these quality filters are exported into individual fastq files with the barcode and probe information trimmed and retained in the read description using a customized SAM TAG. For example, "XM:Z:1:ATCGATCGATCGG:A2:A2" indicates that this paired-end read (R1) has a barcode "ATCGATCGATCGG" and extension and ligation probe sequences matching the STAMP probe pair "A2". This information is retained in the alignment files of stamp as well.

The paired-end reads are then aligned to the complete reference genome containing both nuclear DNA (genome assembly GRCh38) and mtDNA (Revised Cambridge Reference Sequence, rCRS) sequences (ftp://ftp.1000genomes.ebi.ac.uk) using "bwa mem" (1):

**bwa mem -L 100, 5 -M genome_reference.fa filtered_R1.fastq filtered_R2.fastq**

"-L 100,5" disables soft clips following the trimmed probe arm sequences in the paired-end reads. These reads are then aligned again to a revised rCRS with the final 120bp copied to the start to accommodate alignment of reads in the D-loop region:

**bwa mem -L 100, 5 -M shifted_mtdna.fa filtered_R1.fastq filtered_R2.fastq**

The paired-end reads that are unmapped, not in proper pairs, or not aligned to the correct chromosome or locations as per the design of EL probe targets (MAPQ<20), are excluded. The paired-end reads from the 46 mtDNA EL probe pairs are marked as "NUMTS" in the alignment file if they are mapped to nDNA in the complete reference genome (MAPQ≥10).

The properly aligned paired-end reads are locally realigned with freebayes (2) and the base qualities are recalibrated with samtools (3):

**bamleftalign -c -f | samtools calmd -EArb**

Based on the attached molecular barcode, the recalibrated paired-end reads are grouped into read families. The sequence of the consensus read is determined for each read family using a Bayesian approach. In brief, the posterior probability of having a nucleotide, such as "A", at a certain position in the consensus read can be represented using the equation below,

$$P(A|\text{all reads}) = \frac{\prod_{i=1}^{n} P(\text{read}_i|A) \times P(A)}{\sum_{NT} \prod_{i=1}^{n} P(\text{read}_i|NT) \times P(NT)}$$

, where $P$(NT) is prior probability and $\prod_{i=1}^{n} P(\text{read}_i|NT)$ is the estimated likelihood, under the assumption that all paired-end reads in a read family are independent. To simplify calculation, we use equal prior probability for all nucleotides. The likelihood of a nucleotide in each read can be approximated by using the base quality score as

$$P(read_i|\text{NT}) = \begin{cases} 1\text{-}10^{-\frac{\text{BAQ}}{10}}, & \text{NT="A"} \\ \frac{1}{3} \times 10^{-\frac{\text{BAQ}}{10}}, & \text{NT}\neq\text{"A"} \end{cases}$$

The nucleotide with the highest posterior probability ($P_{max}$) is used to construct the consensus read, and assign a quality to this nucleotide by using the phred score of its probability as -10log$_{10}$(1-$P_{max}$). The quality scores of the consensus read are rounded to the nearest integers and are stored in a bam file with ASCII characters from 33 to 126. So, the maximum phred quality score of a nucleotide is 93, which is equivalent to an error rate of <$10^{-9}$. Finally, consensus reads are exported as single-end reads, along with their base quality information into a bam file, for each individual sample. Read information such as “NUMTS” and the number of nucleotide mismatches to the rCRS or the major mtDNA sequence of the sample are exported as additional annotations in the alignment file.

**Here is the command to list the arguments of “stamp align”:**

**python stamp.py align --help**

```
usage: stamp align [-options] sample

positional arguments:
  sample                    sample name

optional arguments:
  -h, --help                show this help message and exit
  -v, --version             show program's version number and exit
  -r1 R1 [R1 ...], --read1 R1 [R1 ...]
                            fastq file(s) for read 1
  -r2 R2 [R2 ...], --read2 R2 [R2 ...]
                            fastq file(s) for read 2
  -p PROBE, --probe PROBE
                            the file with the probe information
  -o OUTPATH, --outpath OUTPATH
                            path where to store the temporary alignment files
 --consensus-outpath CONSENSUS_OUTPATH
                            path where to store the consensus alignment files (default: outpath)
  --override                override output files (default: skip exiting output files)
  --genome GENOME           the complete genome reference
  --mtdna MTDNA             the mtDNA reference sequence
  --mtdna-offset MTDNA_OFFSET
                            the position offset used in parsing mtDNA read alignments
  --numts NUMTS             known nuclear mitochondrial DNA segments (HSD file)
  …
```

**Command to perform read alignment and consensus read calling:**

```
#perform alignment and call consensus reads for sample1

python stamp.py align -r1 sample1.R1.fastq -r2 sample1.R2.fastq --genome
GRCh38_full_analysis_set_plus_decoy_hla.fa --mtdna rCRS_chrM_16449-1-16569.fa
--mtdna-offset 120 --probe stamp_el_probe.txt --numts nmuts_sites_hg38.rCRS.hsd --outpath
bam --consensus-outpath consensus.realign.recal/bam sample1
```

**Important arguments**

- **-r1, -r2:** the R1 and R2 files containing paired-end reads. Stamp can read and combine multiple R1 and R2 files of a sample.
- **--genome:** the complete human genome sequence in FASTA format with bwa index.
- **--mtdna:** the mtDNA sequence in FASTA format with bwa index.
- **--mtdna-offset:** the default offset in relation to the rCRS is 120bp.
- **--numts:** the file with a collection of known NUMTS sequences in the reference genome in HSD format (http://haplogrep.uibk.ac.at/blog/tag/hsd/).
- **--probe**: the file containing the EL probe information (Table 1).
- **--output**: the output directory for the alignment files of the paired-end reads.
- **--output-consensus**: the output directory for the alignment files of the consensus reads.

**Output files**

- **bam/{sample_prefix}_R(1/2).fastq.gz**: the fastq files with barcode and probe information trimmed and stored as an annotation in read description (XM:Z:1:barcode:probe1:probe2).
- **consensus.realign.recal/bam/{sample_prefix}.mtdna.sorted.realign.recal.bam:** the alignment file of paired-end reads generated after base recalibration.
- **consensus.realign.recal/bam/{sample_prefix}.mtdna.consensus.bam:** the alignment file of consensus reads. Consensus reads are output as single-end reads constructed from paired-end reads with the same barcode (read family). Read names are assigned using that of the first paired-end read in a read family. The number of paired-end reads and the barcode is recorded in the SAM TAGs "XF" and "XM" of the consensus read.

**B2. Post-alignment processing and base summarization**

In the "pileup" module, stamp reads the alignment file of the consensus reads (.consensus.bam) and filters the consensus reads according to a list of quality control criteria specified in the arguments. The bases in the retained consensus reads are summarized using samtools to produce a pileup file against mtDNA:

**samtools mpileup -q 20 -Q 0 -B -d 500000 -f mtdna.fa**

The coordinates are further adjusted in the pileup file to the original coordinates of the rCRS.

**Here is the command to list the arguments of "stamp pileup":**

```
python stamp.py pileup --help

    usage: stamp pileup [-options] sample

    positional arguments:
      sample                sample name

    optional arguments:
      -h, --help            show this help message and exit
      -v, --version         show program's version number and exit
      -a ALIGNMENT_FILE [ALIGNMENT_FILE ...], --alignment ALIGNMENT_FILE
    [ALIGNMENT_FILE ...]    the consensus read alignment file(s) output from align
      -p PROBE, --probe PROBE
                            the file with the probe information
      -o OUTPATH, --outpath OUTPATH
                            path where to store the pileup file
      -z, --gzip            compress the pileup file with gzip
      --mtdna MTDNA         the mtDNA reference sequence
      --mtdna-offset MTDNA_OFFSET
                            the position offset used in parsing mtDNA read alignments
      --fs-min FS_MIN       the minimum read family size of consensus reads
      --fs-max FS_MAX       the maximum read family size of consensus reads
      --nm-max NM_MAX       the maximum number of mismatches of consensus reads to the major
                            mtDNA sequence in the coding region
      --nm-max-dloop NM_MAX_DLOOP
                            the maximum number of mismatches of consensus reads to the major
                            mtDNA sequence in the Dloop region
      --tag-excl TAG_EXCL   exclude consensus reads with the tags specified
      --tag-incl TAG_INCL   include consensus reads with the tags specified
      --numts-excl NUMTS    exclude reads from nuclear mitochondrial DNA segments
                            specified by the HSD file
      …
```

**Command to filter consensus reads and summarize base information:**

```
#exclude consensus reads with excessive mismatches or NUMTS annotations
python stamp.py pileup -a sample1.mtdna.consensus.bam --mtdna rCRS_chrM_16449-1-16569.fa
--mtdna-offset 120 --probe stamp_el_probe.txt --numts-excl nmuts_sites_hg38.rCRS.hsd
--nm-max 5 --nm-max-dloop 8 --tag-excl NUMTS,EXMISMATCH
-o consensus.realign.recal/var sample1
```

```
#continued
#only retain consensus reads constructed from duplicate paired-end reads
python stamp.py pileup -a sample1.mtdna.consensus.bam --mtdna rCRS_chrM_16449-1-16569.fa
--mtdna-offset 120 --probe stamp_el_probe.txt --numts-excl nmuts_sites_hg38.rCRS.hsd
--nm-max 5 --nm-max-dloop 8 --fs-min 2 --tag-excl NUMTS,EXMISMATCH
-o consensus.realign.recal/var.f2 sample1
```

##### Important arguments

- **-a**: the consensus read alignment file(s) that are output from align; stamp can read and combine multiple alignment files of a sample.
- **-o**: the file path where to store the pileup file.
- **--mtdna:** the mtDNA sequence in FASTA format with bwa index.
- **--mtdna-offset:** the default offset in relation to the rCRS is 120bp.
- **--numts-excl:** the file with a collection of NUMTS sequences in HSD format.
- **--probe**: the file containing the EL probe information (Table 1).
- **--tag-excl/--tag-incl:** filter out or retain consensus reads using quality information in SAM TAG "XQ"

##### Output file

- **consensus.realign.recal/var/{sample_prefix}.pileup:** the resulting pileup file, which is compressed if "-z" is turned on.

##### B3. mtDNA variant detection

In the "scan" module, stamp reads the pileup file to call mtDNA homoplasmies and heteroplasmies. In contrast to the "align" and "pileup" modules, "stamp scan" can use the pileup files produced from other sequencing methods and computational tools.

##### Here is the command to list the arguments of "stamp scan":

```
python stamp.py scan --help

    usage: stamp scan [-options] input output

    positional arguments:
      input                 a mpileup file or a batch file
      output                the prefix of output files

    optional arguments:
      -h, --help            show this help message and exit
      -v, --version         show program's version number and exit
```

```
#continued

  --mind MIN_DEPTH          the minimum read depth to call variants
  --mind-fwd MIN_DEPTH_FWD
                            the minimum read depth on the forward strand to call variants
  --mind-rev MIN_DEPTH_REV
                            the minimum read depth on the reverse strand to call variants
  --minh MIN_MINOR_DEPTH
                            the minimum number of minor alleles to call heteroplasmies
  --minh-fwd MIN_MINOR_DEPTH_FWD
                            the minimum number of minor alleles on the forward strand to call
                            heteroplasmies
  --minh-rev MIN_MINOR_DEPTH_REV
                            the minimum number of minor alleles on the reverse strand to call
                            heteroplasmies

  --min-het MIN_HET_FREQ
                            the minimum minor allele fraction of heteroplasmies
  --min-qual MIN_QUAL       the minimum base quality
  --max-qual MAX_QUAL       the maximum base quality
  --min-qual-rate MIN_QUAL_RATE
                            the minimum proportion of bases passing the quality filter(s)
  --mle                     use maximum likelihood estimation to compute variant
                            quality for heteroplasmies
  --family FAMILY           family name
  --sample SAMPLE [SAMPLE ...]
                            sample columns to process in the mpileup file
  --name NAME [NAME ...]
                            the corresponding names of the samples to process
  --batch                   proceed in the batch mode (read arguments from the batch file)
  --all-sites               output information for all sites instead of only variant sites
  ....
```

#### Command to call mtDNA variants:

```
#call mtDNA homoplasmies and heteroplasmies of VAF≥1%

python stamp.py scan --name sample1   --mind 100 --mindh 500 --min-qual 30 --minh 5 --min-het 0.01 --mle sample1.mtdna.consensus.adj.pileup.gz consensus.realign.recal/var/sample1.q30
```

**Important arguments**

- **input:** the input mpileup/pileup file; when "--batch" is turned on, "stamp scan" will read the arguments and the path of the pileup file from each line in the batch file provided.
- **--sample:** indicates the order of the column(s) to process in the mpileup file.
- **--name:** the name(s) of the sample(s) in the mpileup file.
- **--family:** a family identifier of the sample(s).

**Output file**

**consensus.realign.recal/var/sample1.q30.var**: a text file (in TSV format) which contains information on the variants identified; it contains information for all mtDNA sites if "--all-sites" is turned on.

#### B4. mtDNA variant annotation

"stamp annot" adds function annotation to the variants in the file generated from "stamp var" using the provided mtDNA annotation database. It also converts this text file into other formats such as ped and hsd which can be read by other bioinformatics tools for mtDNA variant analysis.

**Here is the command to list the arguments of "stamp annot":**

```
python stamp.py annot --help

    usage: stamp annot [-options] input output

    positional arguments:
      input                   the variant file output from scan
      output                  the prefix of output files

    optional arguments:
      -h, --help              show this help message and exit
      -v, --version           show program's version number and exit
      --exclude EXCLUDE       exclude mtDNA sites from analysis
      --remove REMOVE         remove families from analysis
      --keep KEEP             keep only the families for analysis
      --depth DEPTH           the minimum read depth of all variants
      --depth-min DEPTH_MIN
                              the minimum read depth of heteroplasmies
      --hq-min HQ_MIN         the minimum rate of high-quality reads of heteroplasmies
      --llr-min LLR_MIN       the minimum quality score of heteroplasmies
```

```
#continued
    --sbias-min SBIAS_MIN
                          the minimum P value for strand bias analysis of heteroplasmies
    --frac-min FRAC_MIN   the minimum minor allele fraction of heteroplasmies
    --dev-frac-min DEV_FRAC_MIN
                          the minimum variant allele fraction of homoplasmies
    --annotate ANNOTATE   annotate variants according to the file specified
    --output-ped          output the variants detected to a ped file
    --output-hsd          output the major alleles to a hsd file
    --output-minor-hsd    output the minor alleles to a hsd file
   …
```

**Command to annotate mtDNA variants:**

```
#annotate mtDNA variants for all samples
python stamp.py annot --exclude low_quality_sites.txt --remove low_quality_samples.txt --frac-min 0.01 --output-ped --output-hsd --annotate annovar.tsv all_samples.q30.var.combined all_samples.q30.var
```

**Important arguments**

- **input:** the var file from stamp scan, or the file obtained by merging multiple var files.
- **--exclude; --remove; --keep:** the text file containing a list of mtDNA sites or sample IDs to be excluded or included.
- **---output-ped; --output-hsd; --output-hsd-minor:** convert information in the var file to a ped or hsd file.
- **--annotate:** a text file with annotation information for mtDNA.

**An example of the annotation file**

| pos | ref | alt | id | gene | function | variability | freq |
|---|---|---|---|---|---|---|---|
| 1 | G | A | G1A | D-loop | D-loop | 0 | 0 |
| 1 | G | C | G1C | D-loop | D-loop | 0 | 0 |
| 1 | G | T | G1T | D-loop | D-loop | 0 | 0 |
| 2 | A | C | A2C | D-loop | D-loop | 0 | 0 |
| 2 | A | G | A2G | D-loop | D-loop | 0 | 0 |
| …. | | | | | | | |

The values in the id column of the annotation table are used to link information between variants in the annotation table and variants in the var file. The last two columns indicate the variability and frequencies of the variants in the general population obtained from HmtDB (4).

**Output files**

**all_samples.q30.var.qc.annot**: a text file (in TSV format) contains information on the variants passing all quality filters and the related mtDNA annotation. The status column in this file indicates the type of the variants: homoplasmy, heteroplasmy, and possible heteroplasmy. “Possible heteroplasmy” indicates a variant which has VAF great than the provided cutoff but does not pass other quality control filters.

**all_samples.q30.var.qc.hsd :** the hsd file recording the major allele information different from the reference mtDNA sequence.

**all_samples.q30.var.qc.{tped/map/fam}:** the plink files recording the major allele information different from the reference mtDNA sequence.

#### Supplementary Figures

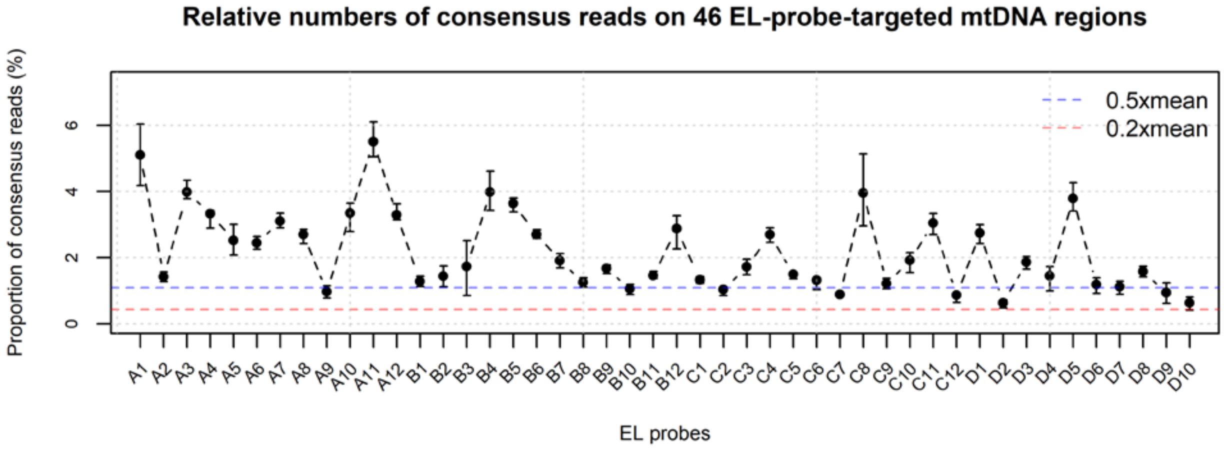


**Figure S1. Relative numbers of consensus reads from the 46 EL-probe-targeted regions in mtDNA.** Each dot refers to the average proportion of consensus reads from each of 46 mtDNA EL probes estimated using reads from 182 lymphoblast samples of REGISTRY. Error bars represent the interquartile range. The blue and red dashed lines indicate 50% and 20% of the mean depth of coverage of consensus reads on mtDNA.

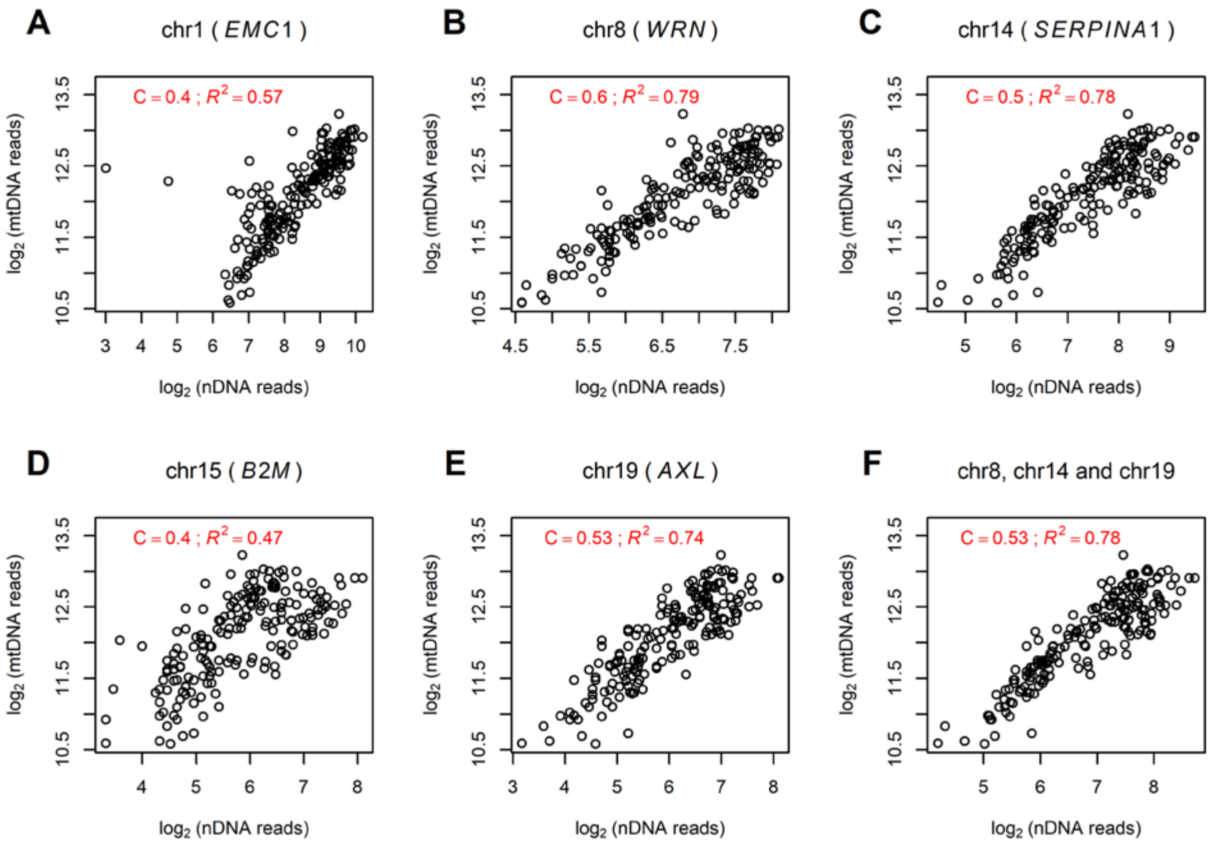


**Figure S2. Correlations of the numbers of consensus reads from mtDNA EL probes and nDNA EL probes.** The results were estimated based on consensus reads from 182 lymphoblast samples of REGISTRY. The number of consensus reads from mtDNA EL probes was averaged over 18 EL probes (A5-A8, B2, B6, B7, B9, C1-C5, C7, C9, C12, D1, and D5). Relationships of mtDNA consensus reads with consensus reads from (**A-E**) each of the 5 nDNA EL probes and (**F**) reads from EL probes targeting chromosomes 8, 14, and 19 were assessed by using linear regression. The number of consensus reads was log transformed with a base of 2. The coefficients C and $R^2$ obtained from each of the regression analyses were indicated in the figure.

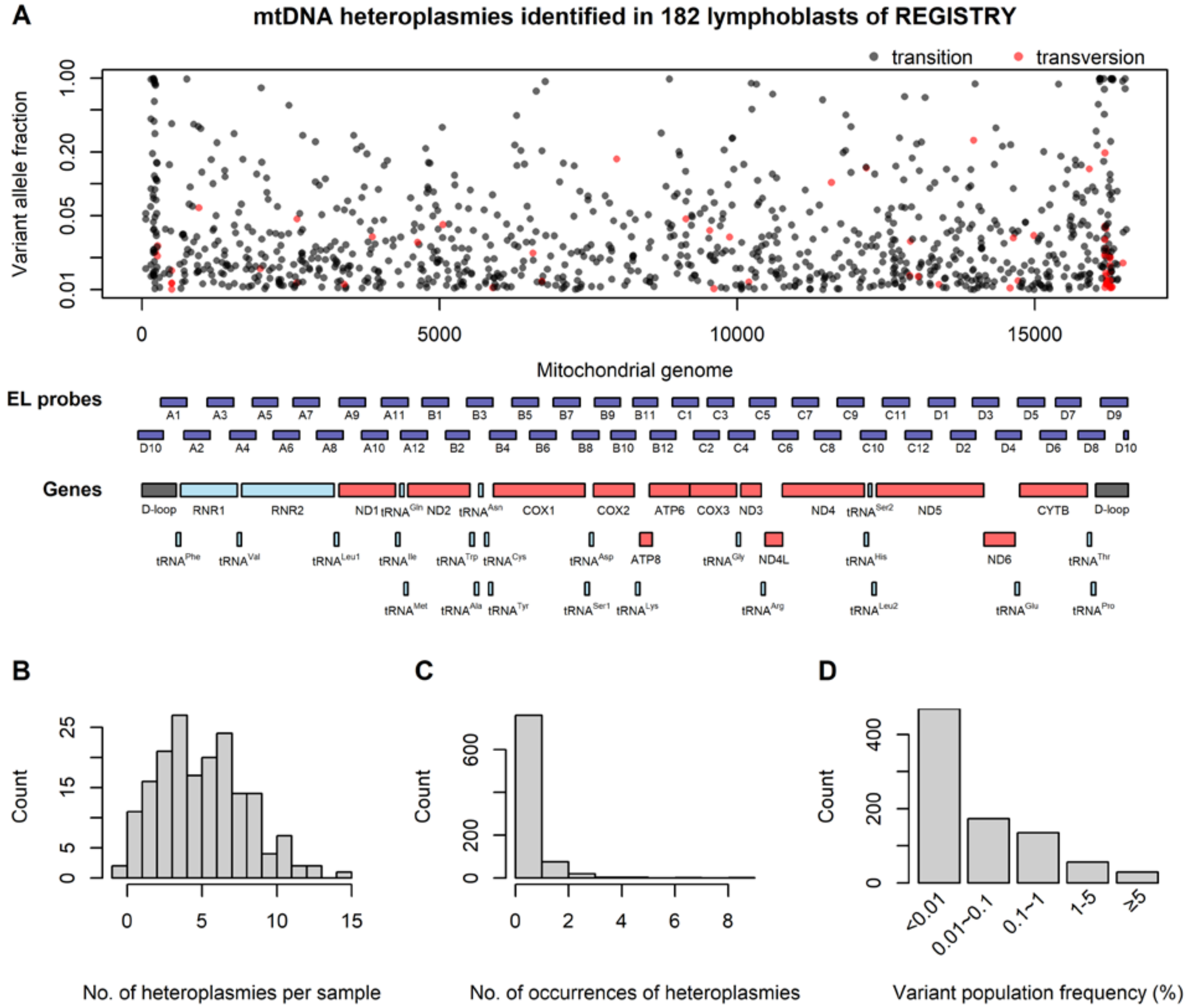


**Figure S3. Summary of heteroplasmies identified in 182 lymphoblast samples of REGISTRY.** (**A**) Positions in mtDNA and variant allele fractions of the 1007 mtDNA heteroplasmies of VAF≥1%. The variant allele fractions are shown on a log scale indicated by the values on the y axis. Heteroplasmies that lead to transition and transversion base changes were depicted in black and red, respectively. The middle panel shows the locations of the 46 EL probes pairs of STAMP as well as the locations of RNA-coding genes (light blue), protein-coding genes (red) and D-loop region (black) of human mtDNA. (**B**) Histogram of the incidence of heteroplasmies per sample. (**C**) Histogram of the occurrence of the 862 heteroplasmic sites. The first bar indicates the number of heteroplasmies unique to one of the 182 lymphoblast samples of REGISTRY. (**D**) Population frequencies of the 862 heteroplasmic sites. Each bar represents the number of heteroplasmic sites with population frequencies in each of the following five categories (<0.01%, 0.01%~0.1%, 0.1%~1%, 1%~5% and ≥5%). Population frequencies of the variants at these sites were obtained from HmtDB.(4)

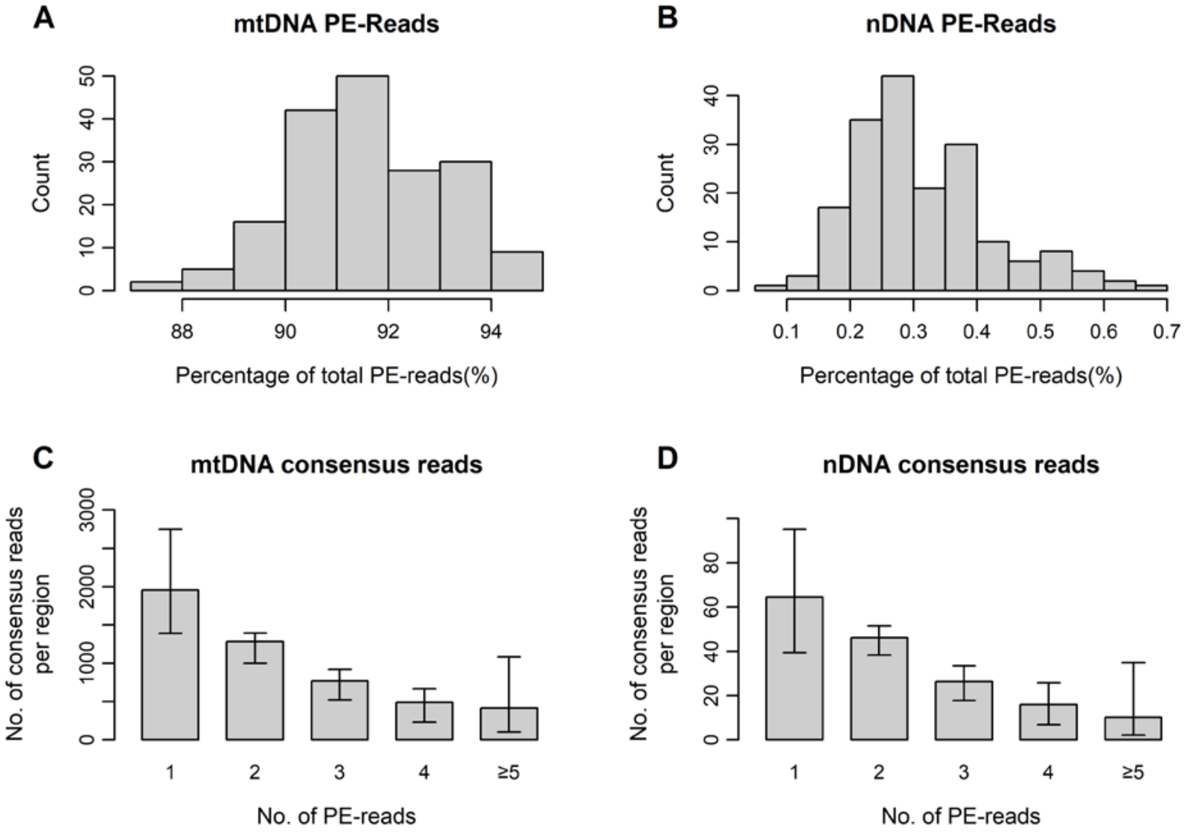


**Figure S4. Mapping rate and duplication rate of paired-end reads in STAMP.** The results were estimated based on paired-end (PE) reads and consensus reads from 182 lymphoblast samples of REGISTRY. The distributions of the percentage of paired-end reads mapped to the EL-probe-targeted regions in (**A**) mtDNA and (**B**) nDNA were depicted using histograms. The distributions of the average numbers of consensus reads constructed from increasing numbers of paired-end reads were depicted in (**C**) for mtDNA-probe-targeted regions and in (**D**) for nDNA-probe-targeted regions. Error bars in (**C**) and (**D**) represent the interquartile range.

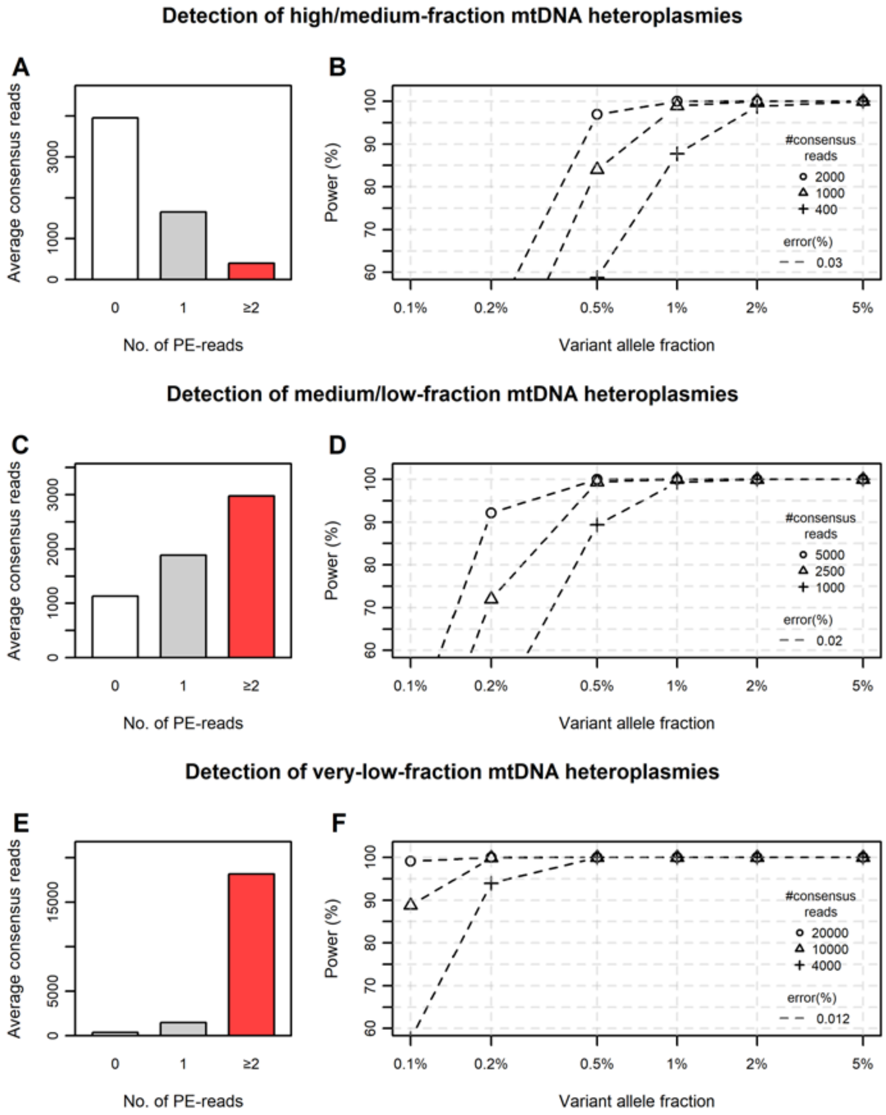


**Figure S5. Power of detecting mtDNA heteroplasmies by using STAMP.** The average numbers of consensus reads with and without duplication in **A**, **C** and **E** were estimated based on a *Poisson* distribution with the numbers of consensus reads and paired-end reads provided in Table S8. The corresponding error rates of STAMP were computed based on the proportions of variant alleles per base in the consensus reads constructed with and without duplication shown in Figure 4D. The statistical power to discriminate real heteroplasmies of varying VAFs from errors, at 16569 sites of mtDNA, was estimated using one-tailed power calculation for one sample proportion. The statistical power was computed with the error rates and the numbers of consensus reads indicated in the legends of panels **B**, **D** and **F**. The statistical power with 50% and 20% of the average number of consensus reads was also depicted for the low-coverage regions in mtDNA. The related results are shown in **A** and **B** for detecting high/medium-fraction heteroplasmies, in **C** and **D** for detecting medium/low-fraction heteroplasmies, and in **E** and **F** for detecting very-low-fraction heteroplasmies.

#### Supplementary Table Titles

**Table S1.** Information of the EL Probes and their target regions used in STAMP.

**Table S2.** Nuclear DNA regions having high sequence similarity to the EL-probe-targeted mtDNA regions.

**Table S3.** Summary of STAMP sequencing in the sample mixtures.

**Table S4.** mtDNA variants identified in the sample mixtures.

**Table S5.** Quality of the mtDNA variants identified in the sample mixtures.

**Table S6.** mtDNA heteroplasmies detected in STAMP replicates of REGISTRY samples.

**Table S7.** Experimental costs in STAMP library preparation.

**Table S8.** Examples of STAMP applications.
